# The parasitic plant *Phtheirospermum japonicum* suppresses host immunity

**DOI:** 10.64898/2026.01.20.700512

**Authors:** Durga Prasad Naik Bhukya, Thomas Spallek

## Abstract

The parasitic plant *Phtheirospermum japonicum* invades the host root xylem to extract nutrients and water. We show that this invasion triggers a phosphate-starvation response in its host, *Arabidopsis thaliana*, causing systemic suppression of immunity, including defence responses against microbial pathogens. Consequently, plants infested by xylem-feeding parasitic plants were more susceptible to secondary infections, underscoring the need to consider multitrophic interactions when improving crop resilience to parasitic infestation.

## Main

Parasitic plants of the Orobanchaceae family cause billions of dollars in crop losses by reducing host growth and yield^1,2^. Although progress has been made in understanding how these parasites recognize hosts and develop haustoria, their infection and feeding organs, little is known about how parasitism impacts host physiology and interactions with other pathogens^3-7^. To experimentally test the effect of parasitism on the host, we quantified individual immune responses in *Arabidopsis thaliana* (*Arabidopsis*) parasitized by *Phtheirospermum japonicum* (*Phtheirospermum*) and compared them to immune responses in unparasitized plants.

*Arabidopsis* parasitized by *Phtheirospermum* developed smaller rosettes (Fig. 1a, Extended Data Fig. 1a) and exhibited significant reductions in fresh and dry weight (Extended Data Fig. 1c,d). *Phtheirospermum* grew taller and accumulated more biomass when parasitizing *Arabidopsis*, demonstrating that key phenotypes of parasitic interactions observed in crops are captured in our experimental model (Fig. 1a, Extended Data Fig. 1b-d). Next, we challenged parasitized and non-parasitized *Arabidopsis* with flg22, a well-studied elicitor of plant immune responses derived from bacterial flagellin^8^. Reactive oxygen species (ROS) production triggered by flg22 was significantly lower in parasitized compared to non-parasitized plants (Fig. 1b and Extended Data Fig. 2a). Similarly, flg22-induced phosphorylation of mitogen-activated protein kinases (MAPKs) was reduced in parasitized plants (Fig. 1d). In these assays, flg22 receptor *fls2* and co-receptor *bak1-5* mutants served as controls for non-elicitor-specific responses (Fig. 1b,c and Extended Data Fig. 2a). A reduction in immune responses was also observed in host plants treated with chitin, a major component of fungal cell walls (Fig. 1c,e and Extended Data Fig. 2b). To better understand how parasitism affects flg22-triggered immunity, we performed RNA-seq on RNA isolated from *Arabidopsis* leaves that had been infiltrated with or without flg22. Flg22 triggered a substantial transcriptional reprogramming in *Arabidopsis*, which was overall less pronounced in parasitized plants (Fig. 1f and Extended Data Fig. 4). RNAs, which were less abundant in parasitized hosts than in non-parasitized controls, included transcripts from genes along the entire FLS2 signalling pathway (Fig. 1f). Lower-expressed genes included *FLS2* and its co-receptor *BAK1*, downstream components such as *RbohD*, receptor-like cytoplasmic kinases, MAPKs, and defence response genes including *FRK1, WRKY* transcription factors, and salicylic acid pathway components, indicating a broad suppression of immune responses in parasitized plants.

**Fig. 1.**
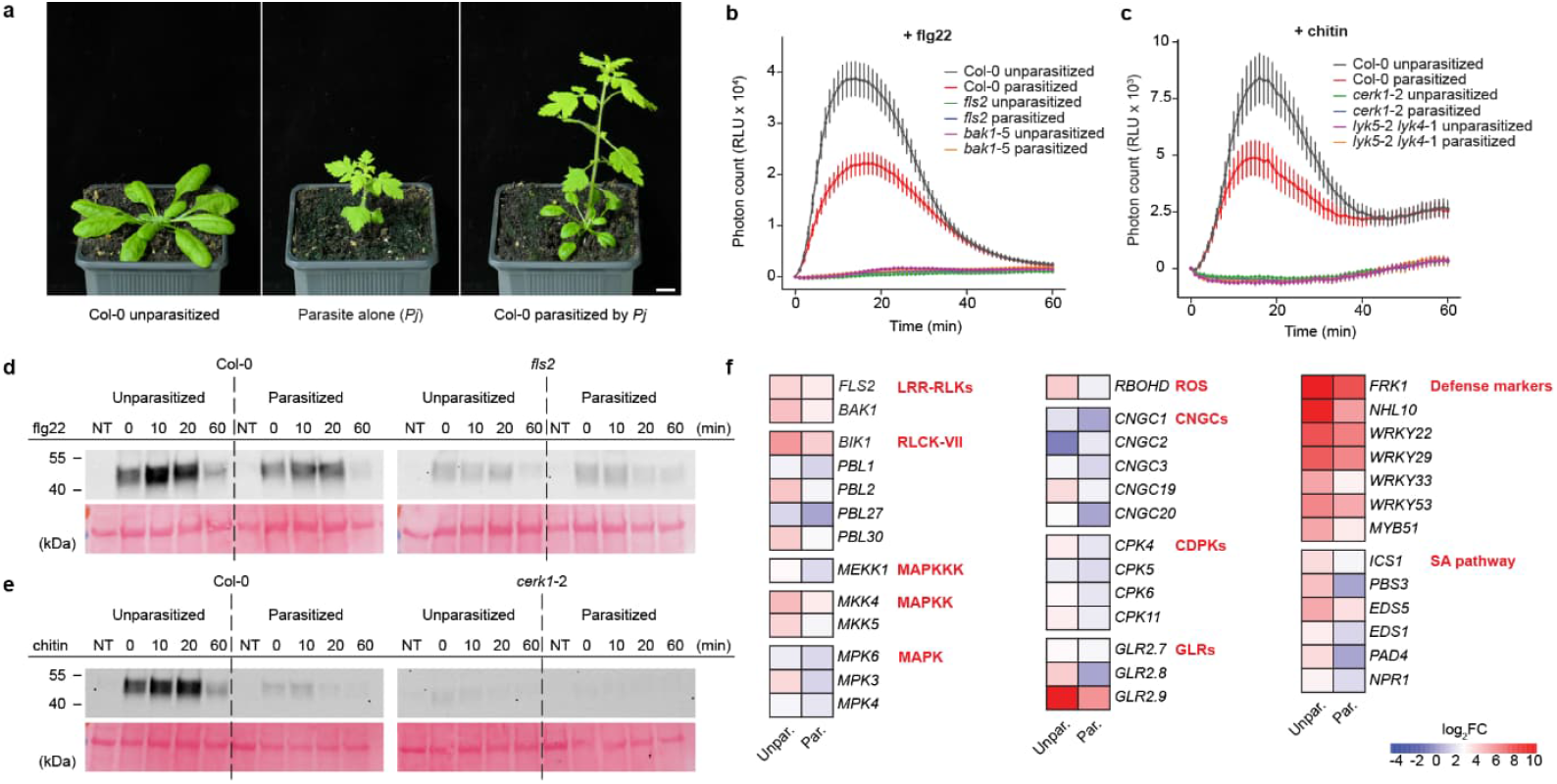
*Phtheirospermum* parasitism suppresses morphological traits, and early and late PTI signalling in *Arabidopsis* Col-0 plants. **a**, Representative images of an unparasitized *Arabidopsis* (left), a non-parasitizing *Phtheirospermum* (middle), and the *Arabidopsis* parasitized by *Phtheirospermum* at 30-day-post parasitization. Scale bars, 1 cm. **b, c**, ROS burst kinetics of unparasitized and parasitized *Arabidopsis* wild-type (WT) Col-0, flg22-receptor mutant *fls2*, and co-receptor mutant *bak1*-5 leaf discs after treatment with 100 nM flg22 **(b)** and *Arabidopsis* WT Col-0, chitin-receptor mutant *cerk1*-2, and co-receptor mutant *lyk5*-2 *lyk4*-1 leaf discs after treatment with 25 µg/mL chitin **(c).** RLU, relative light units. **d, e**, MAPK activation in unparasitized and parasitized *Arabidopsis* WT Col-0, and flg22-receptor mutant *fls2* after treatment with 100 nM flg22 **(d)** and *Arabidopsis* WT Col-0, chitin-receptor mutant *cerk1*-2 after treatment with 25 µg/mL chitin **(e)**; uncropped blots in Extended Data Fig. 3. NT, non-treated. Ponceau S staining is shown as an equal loading control. **f**, Heatmaps showing the expression level of representative genes across key PTI signalling and salicylic acid (SA) pathway in unparasitized and parasitized *Arabidopsis* Col-0 plants at 60 min after treatment with 100 nM flg22. Red indicates transcriptional upregulation and blue indicates downregulation of genes. Data represent log_2_-transformed fold changes relative to untreated unparasitized and parasitized *Arabidopsis* Col-0 plants.

**Fig. 2.**
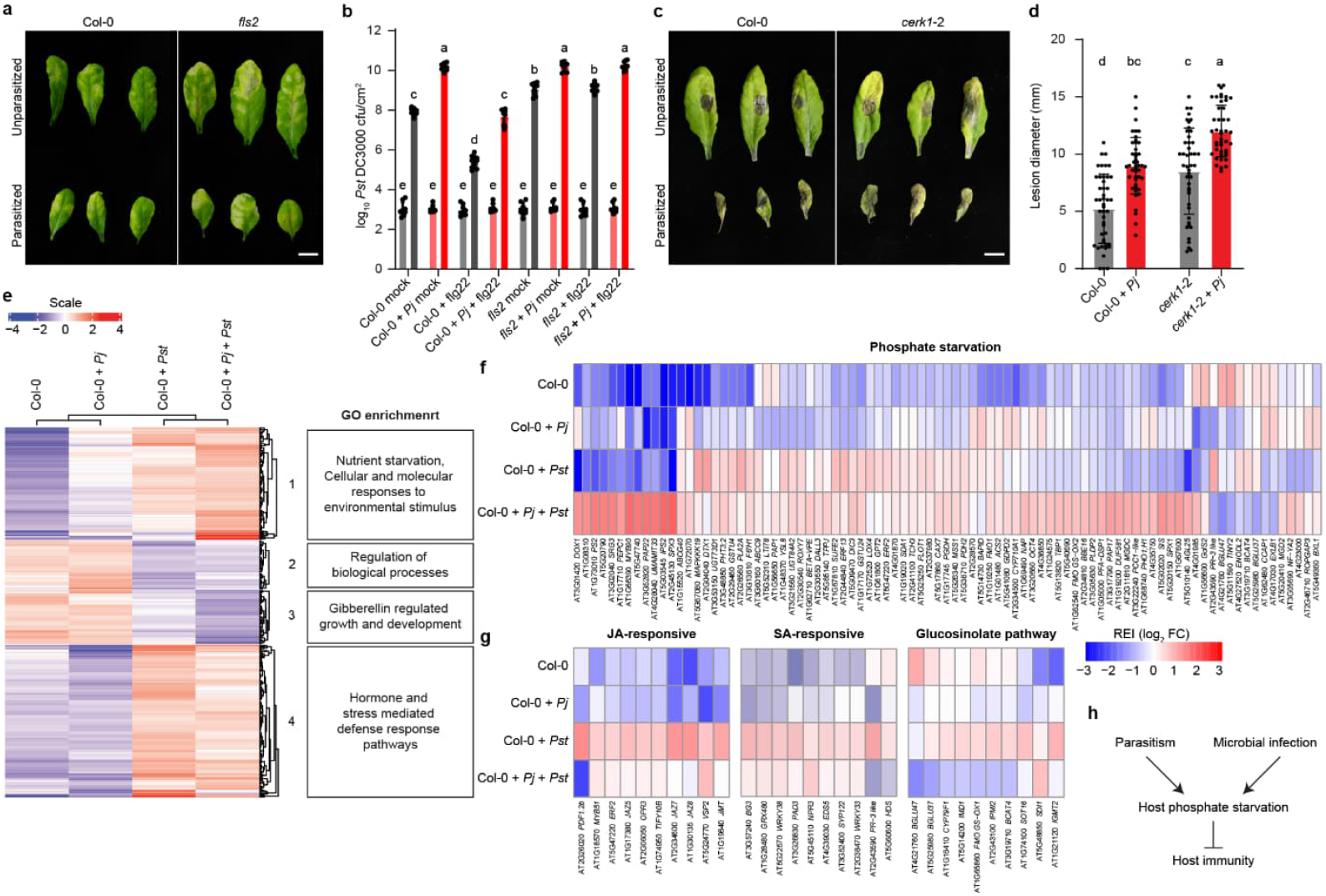
*Phtheirospermum* parasitism increases pathogen susceptibility through nutrient starvation and immune suppression in *Arabidopsis* Col-0 plants. **a**, Disease symptoms of unparasitized and parasitized *Arabidopsis* WT Col-0, and flg22-receptor mutant *fls2* plants at 72 hours post-infection (hpi) with 1 × 10^5^ CFU/mL *Pst* DC3000. Scale bars, 1 cm. **b**, Quantification of bacterial growth in leaves of unparasitized and parasitized *Arabidopsis* WT Col-0, and flg22-receptor mutant *fls2* plants. Leaves were pre-treated with 1 µM flg22 or sterile ddH_2_O (mock) 24 h prior to inoculation with 1 × 10^5^ CFU/mL *Pst* DC3000, and bacterial populations were quantified at 3 hpi and 72 hpi. **c**, Disease symptoms of unparasitized and parasitized *Arabidopsis* WT Col-0, and chitin-receptor mutant *cerk1*-2 plants at 3 dpi with 5 × 10^4^ spores/mL of *Botrytis cinerea*. Scale bars, 1 cm. **d**, Quantification of fungal growth in leaves of unparasitized and parasitized *Arabidopsis* WT Col-0, and chitin-receptor mutant *cerk1*-2 plants. Leaves were drop-inoculated with 5 × 10^4^ spores/mL of *Botrytis cinerea*. **b, d**, Data represent mean +/-SD of n = 3 independent biological replicates. Different letters above the bars indicate statistically significant differences among treatments (one-way ANOVA followed by Tukey’s HSD post hoc test, P < 0.05). **e**, Hierarchical clustering of differentially expressed genes in mock- and *Pst* DC3000-infected (24 hpi) unparasitized and parasitized *Arabidopsis* WT Col-0 plants, identified using DESeq2 (|log_2_FC| ≥ 1, FDR < 0.05). Transcript abundance is displayed as a z-score colour scale calculated from normalized counts, with red indicating higher and blue indicating lower expression relative to the mean across samples. Gene Ontology (GO) enrichment terms are annotated for the expression clusters, with enrichment assessed against the full transcriptome background. **f, g**, Heatmap showing the relative expression profiles of genes involved in the phosphate starvation response **(f)** and jasmonic acid (JA) responsive, salicylic acid (SA) responsive and glucosinolate pathway **(g)** in mock- and *Pst* DC3000-infected (24 hpi) unparasitized and parasitized *Arabidopsis* WT Col-0 plants. Expression values represent the log_2_ fold change of each gene relative to its mean expression across all conditions. REI, relative expression index. Red, higher expression; blue, lower expression. **h**, Schematic of the effect of *Phtheirospermum* parasitism on *Arabidopsis* immunity against microbial pathogens.

Next, we asked whether reduced immune responsiveness to flg22 and chitin translates into altered susceptibility to microbial pathogens. To test this, we syringe-infiltrated leaves of *Arabidopsis* wild-type (Col-0) and *fls2* mutant plants with virulent *Pseudomonas syringae* pv. *tomato* DC3000 (*Pst*). Col-0 developed more severe disease symptoms when parasitized than when unparasitized, phenocopying the hypersusceptible *fls2* mutant (Fig. 2a). In separate experiments, plants were pre-infiltrated one day earlier with either flg22 or a buffer control, and bacteria were re-isolated at 3 and 72 hours after infiltration. While comparable *Pst* titres were recovered across all conditions at 3 hours, parasitized plants showed markedly greater bacterial growth at 72 hours (Fig. 2b). Flg22 infiltration reduced susceptibility to *Pst* in both unparasitized and parasitized Col-0 plants, but not in *fls2* mutants (Fig. 2b). However, flg22-induced resistance was substantially lower in parasitized hosts, with parasitized, flg22-pretreated Col-0 plants showing similar *Pst* growth as Col-0 plants that were not pre-treated with flg22. Hyper-susceptibility of parasitized hosts was also observed in infections with the fungal pathogen *Botrytis cinerea*. Leaves of parasitized wild-type plants were nearly completely necrotic and developed significantly larger lesions, resembling the hypersusceptible *lyk5-2 lyk4-1* double mutants (Fig. 2c,d). We then compared the transcriptomes of parasitized *Arabidopsis* shoots with those of unparasitized plants 24 hours after infiltration with *Pst*. RNA-seq analysis identified 402 differentially expressed genes (DEGs) across the following conditions: unparasitized/mock-infiltrated, parasitized/mock-infiltrated, unparasitized/*Pst*-infected, and parasitized/*Pst*-infected *Arabidopsis* plants (Fig. 2e and Extended Data Fig. 5). These DEGs were grouped into four clusters. Cluster 1 comprised DEGs up-regulated by *Pst* and showing even higher expression in parasitized plants. Gene Ontology (GO) terms associated with this cluster were significantly enriched for responses to environmental stimuli related to nutrient scarcity. Clusters 2 and 3 contained DEGs repressed by *Pst* infection and either less repressed or further repressed in parasitized plants. These genes were linked to growth-related processes. Cluster 4 included DEGs induced by *Pst* that showed lower expression in parasitized plants. These DEGs were enriched for GO terms related to immunity, supporting our earlier observation that parasitized hosts were less capable of mounting an efficient immune response. Notably, cluster 1 included several genes associated with phosphate starvation. Given the well-established role of phosphate starvation in modulating plant immune responses to microbial pathogens and symbionts^9-12^, we examined the expression profiles of phosphate-starvation-responsive genes. Indeed, their expression was markedly elevated in *Pst*-infected, parasitized plants compared to the other three conditions (Fig. 2f). Parasitism and *Pst* infection alone also increased the expression of these genes relative to control plants. This effect was further amplified when plants were exposed to both pathogens simultaneously. Consistent with the known influence of phosphate starvation on plant immunity, genes central to the SA, JA, and glucosinolate pathways were less efficiently induced during co-infection, supporting our initial observation that parasitized plants are impaired in mounting effective defence responses to individual immune elicitors as well as to bacterial and fungal pathogens (Fig. 2f).

In summary, our study highlights a previously overlooked effect of plant parasitism on host immunity. Non-selective uptake of xylem-transported substances by the parasite, as shown by dye-uptake experiments, reduces phosphate availability in host shoots^13^. Given that phosphate starvation attenuates immune responses, this nutrient depletion provides a mechanistic explanation for the compromised immunity observed in *Arabidopsis* parasitized by *Phtheirospermum* (Fig. 2h). Parasitized plants are therefore more vulnerable to secondary infections. Indeed, maize parasitized by *Striga* has been reported to show increased susceptibility to spotted stem borer, *Chilo partellus*^14^. These findings underscore the importance of considering combined stress effects when assessing the impact of parasitic plants on crops.

## Methods

### Plant material and growth conditions

*Phtheirospermum japonicum* (Thunb.) Kanitz ecotype Okayama wild type used as the parasitic plant^15^. The *Arabidopsis thaliana* ecotype Columbia-0 (Col-0), the flg22 receptor mutants *fls2*^8^ *bak1*-5^16^ and the chitin receptor mutants *cerk1*-2^17^ *lyk5*-2 *lyk4*-1^18^ used as host plants. Plant growth conditions were based on a previously published protocol with minor modifications^6^. In brief, *Phtheirospermum* and *Arabidopsis* seeds were surface sterilized in 70% (v/v) ethanol containing 0.05% (v/v) SDS for 10 min on an overhead shaker. After sterilization, seeds were rinsed five times with 100% ethanol, transferred to sterile filter paper, air-dried completely. Sterilized *Phtheirospermum* seeds were sown on 0.8% water agar (Duchefa, Cat# P1001), while *Arabidopsis* seeds were plated on full-strength Murashige & Skoog medium (MS; Duchefa, Cat# M0222) supplemented with 0.8% plant agar (Duchefa, Cat# P1001) and adjusted to pH 5.8. Plates were stratified at 4 °C for 2-3 days and subsequently grown vertically under short-day (SD) conditions (12 h light/12 h dark, 22 °C, 60% relative humidity, 100 µmol m^-2^ s^-1^).

### Parasitization assay

Two-week-old *Arabidopsis* seedlings grown on full-strength MS were placed adjacent to one-week-old *Phtheirospermum* seedlings maintained on water agar. The roots of the *Phtheirospermum* and *Arabidopsis* were gently aligned in direct contact and oriented in the same direction. Plates were positioned vertically and maintained under SD conditions for 7 days, with control *Phtheirospermum* and *Arabidopsis* seedlings grown separately on water agar plates under same conditions. Following this interaction period, *Phtheirospermum*-*Arabidopsis* pairs, along with non-parasitizing *Phtheirospermum* and unparasitized *Arabidopsis* controls, were transferred to steam-sterilized soil pre-saturated once with 0.2% Wuxal Super (Manna, Ammerbuch-Pfäffingen, Germany). Plants were subsequently grown under SD conditions for an additional four weeks to assess parasitism outcomes.

### Microbial pathogen infection assays

*Pseudomonas syringae* pv. *tomato* DC3000 (*Pst* DC3000) was cultured in NYG medium (5 g/L peptone, 3 g/L yeast extract, 20% glycerol, pH 7.0) supplemented with rifampicin (50 μg/mL) and kanamycin (50 μg/mL) at 28 °C to an OD_600_ of ∼0.2. Bacterial cells were harvested by centrifugation (4000 g for 10 min), washed twice with 10 mM MgCl_2_, and adjusted to 10^5^ cfu/mL in 10 mM MgCl_2_. Bacterial suspensions were pressure-infiltrated into the abaxial leaf surface of unparasitized and parasitized *Arabidopsis* plants using a needleless 1-mL syringe.

For priming experiments, leaves were pretreated with 1 µM flg22 or sterile ddH_2_O 24 h prior to *Pst* DC3000 infiltration as previously described^8^. Leaf discs were collected at 3- and 72-hpi using a 4-mm biopsy punch. For each biological replicate, leaves from three independent plants were sampled; eight discs per plant (two discs from each of four infiltrated leaves) were pooled in 500 µL of 10 mM MgCl_2_ and homogenized with a micro-pestle. Serial 10-fold dilutions (10^-1^-10^-7^) of the homogenate were plated (10 µL) on NYG agar containing the appropriate antibiotics. Colony counts were recorded after 2 days of incubation at 28 °C, and bacterial titres were represented as CFU cm^−2^ of leaf tissue.

*Botrytis cinerea* infection was performed according to previously described method^19^, with minor modifications. Spores of *Botrytis cinerea* strain B05.10 were suspended in sterile one-quarter-strength of PBD (3 g/L potato dextrose broth) to a final concentration of 5 × 10^4^ spores/mL and incubated at room temperature for 4 h to induce germination. A 6-μL droplet of the germinated spore suspension was placed onto the surface of leaves from both unparasitized and parasitized *Arabidopsis* plants. For each biological replicate, three leaves from five independent plants were inoculated. Following inoculation, plants are enclosed with a transparent lid to maintain high humidity. Lesion diameters were recorded 3 days-post inoculation using a digital caliper.

### Phenotypic measurements

Phenotypic traits of *Arabidopsis* (unparasitized and parasitized) and *Phtheirospermum* (non-parasitizing (*Pj*) and parasitizing (*Pj* + Col-0)) plants were assessed four weeks after transfer to soil. Rosette width of *Arabidopsis* was measured across the widest diameter of the rosette using a digital caliper. For *Phtheirospermum*, plant height was measured from the root-shoot junction to the shoot apex. For biomass quantification, aboveground tissues were harvested and weighed immediately to obtain fresh weight, then dried at 60 °C for 3 days before determining dry weight. Measurements were performed on three biological replicates per condition, with 15 plants per replicate.

### Measurement of ROS production

To assess ROS accumulation, leaf discs (Ø = 2 mm) were harvested using a standard biopsy punch (Integra LifeSciences, Cat# 15110-20) from unparasitized and parasitized *Arabidopsis* host plants. All the discs were incubated overnight in ddH_2_O. The next day, water was replaced with 190 µL of a solution containing 20 μM luminol L-012 (FUJIFILM Wako Pure Chemical Corporation, Cat# 120-04891) and 2 µg/mL horseradish peroxidase (Carl Roth GmbH, Cat# 6055.1). Light emission was recorded every min for 15 min using a TECAN infinite® M200 plate reader to obtain baseline readings. Subsequently, either 10 µL of 100 nM flg22 (GLPBIO, Cat# GC32202-1) or 25 µg/mL chitin from shrimp cells (Sigma-Aldrich, Cat# C9752) was added, and light emission was monitored every min for 60 min.

### Protein isolation and detection of MAPK phosphorylation

Leaves were harvested from unparasitized and parasitized *Arabidopsis* host plants at 30 days post-parasitization (dpp) and vacuum-infiltrated for 15 min in 20 mL of elicitor solution (either 100 nM flg22 or 25 µg/mL chitin, prepared in ddH_2_O). Following vacuum infiltration, leaves were snap-frozen in liquid nitrogen after 0, 10, 20, and 60 min and stored at –80 °C for subsequent protein extraction and immunoblotting. Leaves from unparasitized and parasitized *Arabidopsis* host plants that were not vacuum infiltrated (Non-treated, NT) were used as controls for infiltration-induced MAPK phosphorylation. Total proteins were extracted using the protein extraction buffer containing 250 mM sucrose, 100 mM HEPES-KOH (pH 7.5), 5% glycerol, 25 mM NaF, 1 mM Na_2_MoO_4_ ·2H_2_O, 50 mM Na_4_P_2_O_7_ ·10H_2_O, 10 mM EDTA, 1 mM DTT, 0.5% Triton X-100 and the Protease Inhibitor Mix P (SERVA, Cat# 39103.03). Protein concentrations were determined using Bradford reagent (Roti®-Quant, Carl Roth, Cat# K015.1) and then normalized to a final concentration of 1 µg/µL using protein extraction buffer supplemented with Protease Inhibitor Mix P. Protein extracts (10 µg) were denatured at 95 °C in 4x SDS loading-dye containing 200 mM Tris-HCl (pH 6.8), 8% SDS, 400 mM DTT, 0.4% Bromophenol blue, 40% glycerol and loaded onto a 8–16% Mini-PROTEAN® TGX™ precast protein gels (Bio-Rad, Cat# 4561103). After separation of proteins on the protein gel, proteins were transferred to a 0.45 µm nitrocellulose membrane (Amersham™ Protran®, Cat# 10600047) and blocked overnight with TBST containing 5% milk. Following blocking, the membrane was washed three times with TBST and incubated with the primary anti-phospho-p44/42 MAPK rabbit monoclonal antibody (1:2000 in 2% milk, Cell Signaling Technology, Cat# 4370S) for 2 h at room temperature (RT) and secondary peroxidase-linked goat anti-rabbit IgG (1:5000 in 5% milk, ThermoFisher Scientific, Cat# 31460) for 1 h at RT with three washes of 10 min each in between the incubations. After washing the membranes, SuperSignal™ West Dura Extended Duration Substrate (Thermo Fisher Scientific, Cat# 34075) was used for chemiluminescent detection with a ChemiDoc (BioRad). To confirm equal protein loading, membranes were stained with Ponceau S (0,1% w/v in 5% acetic acid, Sigma-Aldrich, Cat# P7170), and subsequently rinsed with H_2_O.

### RNA sequencing analysis

To access the flg22-triggered immunity in unparasitized and parasitized *Arabidopsis* Col-0 plants, leaves were collected at 30 dpp and vacuum-infiltrated for 15 min in 20 mL of 100 nM flg22 solution. Samples were harvested 60 min post-vacuum infiltration. To investigate how parasitism influences host immunity, leaves from unparasitized and parasitized *Arabidopsis* Col-0 plants were syringe-infiltrated with *Pseudomonas syringae* pv. *tomato* DC3000 (*Pst* DC3000) 1 x 10^5^ CFU/mL, or with 10 mM MgCl_2_ as a mock treatment. Leaves were collected 24 h after inoculation. Three independent biological replicates were analysed for both datasets. Total RNA was extracted using the RNeasy Plant Mini Kit (Qiagen, Cat# 74904) with on-column DNase treatment, following the manufacturer’s protocol. RNA samples were submitted to GENEWIZ Germany GmbH for poly(A) enrichment, strand-specific library preparation, and paired-end Illumina sequencing (2 x 150 bp). RNA sequencing analysis was performed using the Galaxy Europe platform^20^. Raw reads were aligned to the *Arabidopsis thaliana* TAIR10 reference genome using RNA STAR with default parameters for paired-end data. The genome index was built using AtRTD3 as the reference annotation^21^. Raw gene-level counts generated by featureCounts were filtered to remove low-expression genes across samples, and only reads with a minimum mapping quality score of 10 were included in the analysis. Normalization was carried out using DESeq2’s median-of-ratios method to account for differences in sequencing depth and RNA composition. Differentially expressed genes (DEGs) were identified by comparing different groups, including *Pst*-infected vs. mock-inoculated samples, flg22-treated vs. untreated samples, and parasitized vs. unparasitized samples within each treatment group. Genes were considered differentially expressed if they met the significance criteria of adjusted p-value (FDR) < 0.05 and absolute log2 fold change ≥ 1. Heatmap of DEGs (Fig. 2e) were generated in iDEP v2.01^22^ using DESeq2-normalized counts. Gene expression patterns across samples were visualized through hierarchical and k-means clustering. Genes within each expression cluster were subjected to Gene Ontology (GO) enrichment analysis using the DAVID functional annotation tool^23^. To derive gene expression profiles (Fig. 2f,g), log_2_ expression ratios were calculated between the normalized expression level of each gene under a given condition and the mean expression of that gene across all conditions. This log_2_ ratio is referred to as the Relative Expression Index (REI), which enables comparison of expression dynamics independent of absolute expression levels.

### Phosphate starvation marker gene sets

Phosphate starvation response (PSR) activation was assessed using two published maker gene sets: 193 core PSR genes identified under in vitro phosphate limitation^9^ and 210 PSR genes derived from contrasting soil phosphorus regimes^24^. These two non-redundant sets were used as predefined PSR gene modules. Mapping to our RNA-seq dataset identified 87 shared PSR genes, which were differentially expressed in at least one of the four experimental conditions (unparasitized/parasitized × mock-/*Pst* DC3000-infiltrated). The PSR marker *IPS2*, although not included in the predefined PSR gene modules, was included in the analysis due to its strong induction in parasitized/*Pst* DC3000-infiltrated *Arabidopsis* plants and its established role as a PSR indicator^25^.

## Acknowledgements

We thank Volker Lipka and Markus Albert for sharing *Arabidopsis* lines. We acknowledge Hannes Ruwe for his comments on this work. The work was supported by grants of the German Research Foundation (RTG2172 PRoTECT, 273134146). DB was supported by the GGNB program PRoTECT.

## Author Contribution

DB performed the experiments and analysed the data. TS led the project and, together with DB, designed the study and wrote the manuscript.

## Data Availability

The RNA-Seq data generated in this study have been deposited in the NCBI Sequence Read Archive (SRA) under BioProject ID PRJNA1404945.

---

**Extended Data Fig. 1.**
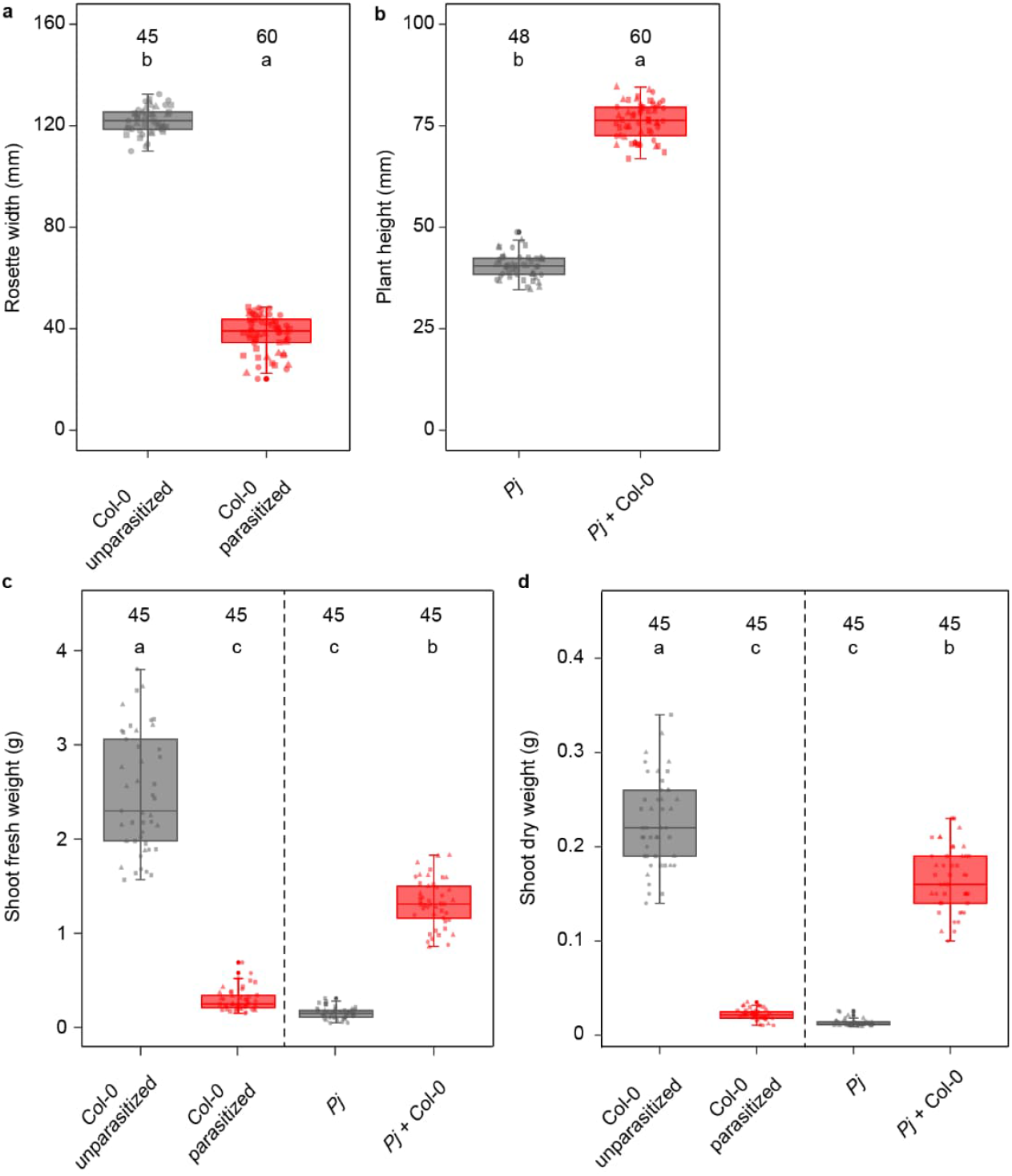
Parasitism alters growth dynamics in *Arabidopsis* Col-0 and *Phtheirospermum*. **a**, Rosette widths of unparasitized and parasitized *Arabidopsis* WT Col-0 plants at 30 days post-parasitization (dpp). **b**, Plant heights of non-parasitizing *Phtheirospermum* (*Pj*) and parasitizing *Pj* (*Pj* + Col-0) at 30 dpp. **c**, Shoot fresh weight biomass of unparasitized and parasitized *Arabidopsis* WT Col-0 plants, as well as *Pj* and *Pj* + Col-0 at 30 dpp. **d**, Shoot dry weight biomass of same genotypes and treatments shown in **(c)**. Different letters indicate statistically significant differences according to one-way ANOVA followed by Tukey’s HSD post hoc test (P < 0.05), based on three independent experiments.

**Extended Data Fig. 2.**
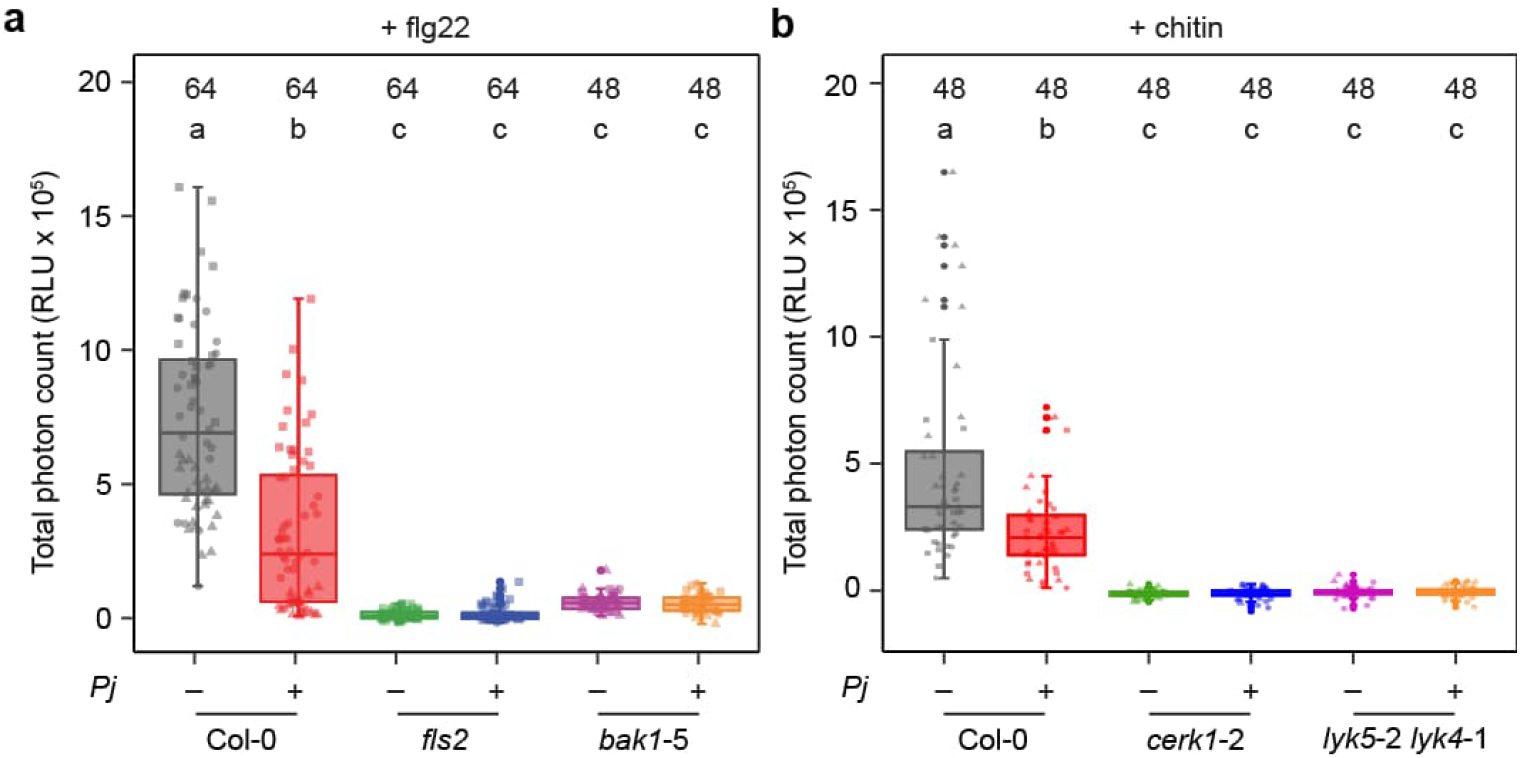
*Phtheirospermum* parasitism suppresses PAMP-triggered ROS production in *Arabidopsis* Col-0 plants. **a, b**, Total ROS production in leaves of unparasitized and parasitized *Arabidopsis* WT Col-0, flg22-receptor mutant *fls2*, and co-receptor mutant *bak1*-5 after treatment with 100 nM flg22 **(a)** and *Arabidopsis* WT Col-0, chitin-receptor mutant *cerk1*-2, and co-receptor mutant *lyk5*-2 *lyk4*-1 leaf discs after treatment with 25 µg/mL chitin **(b)**. Different letters indicate statistically significant differences according to one-way ANOVA followed by Tukey’s HSD post hoc test (P < 0.05), based on three independent experiments.

**Extended Data Fig. 3.**
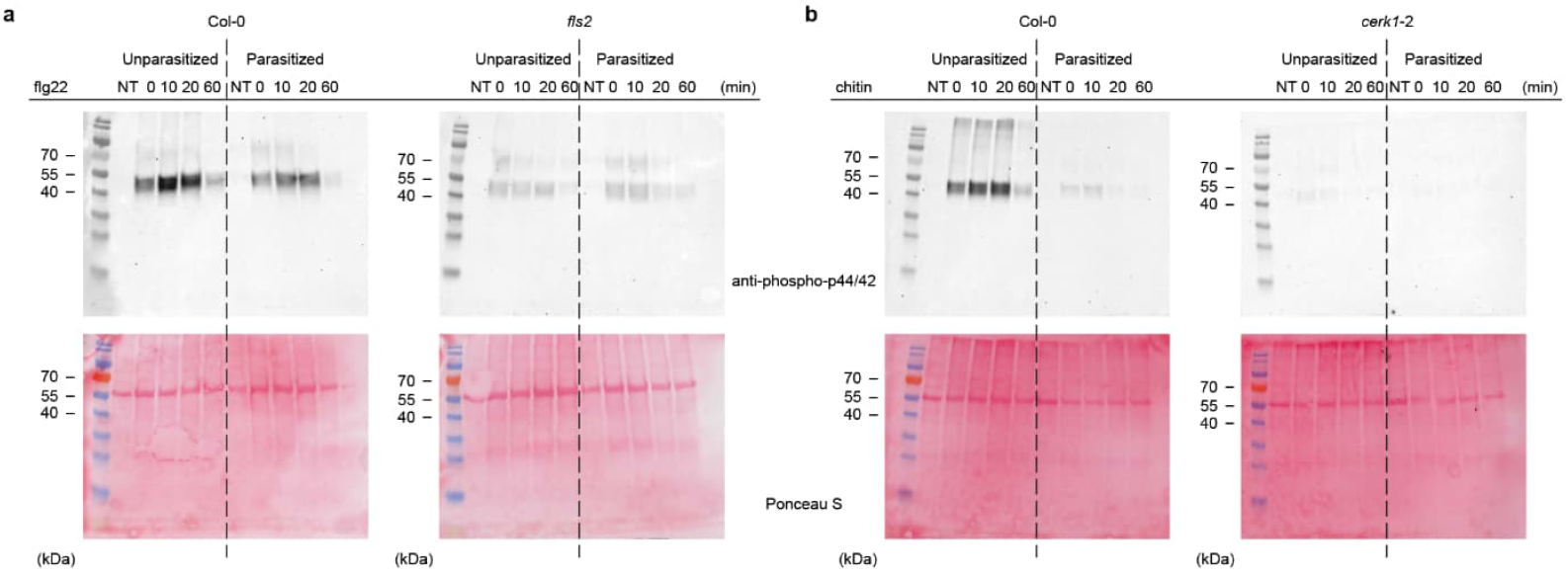
*Phtheirospermum* parasitism suppresses MAPK activation in *Arabidopsis* Col-0 plants. **a, b**, Full, uncropped immunoblots of Fig. 1d,e showing MAPK phosphorylation in leaves of unparasitized and parasitized *Arabidopsis* plants. **a**, WT Col-0 and flg22-receptor mutant *fls2* plants following vacuum infiltration with 100 nM flg22. **b**, WT Col-0 and chitin-receptor mutant *cerk1*-2 plants following vacuum infiltration with 25 µg/mL chitin. NT, non-treated. Phosphorylated MAPKs were detected using anti-phospho-p44/42 antibody. Ponceau S is shown as loading control.

**Extended Data Fig. 4.**
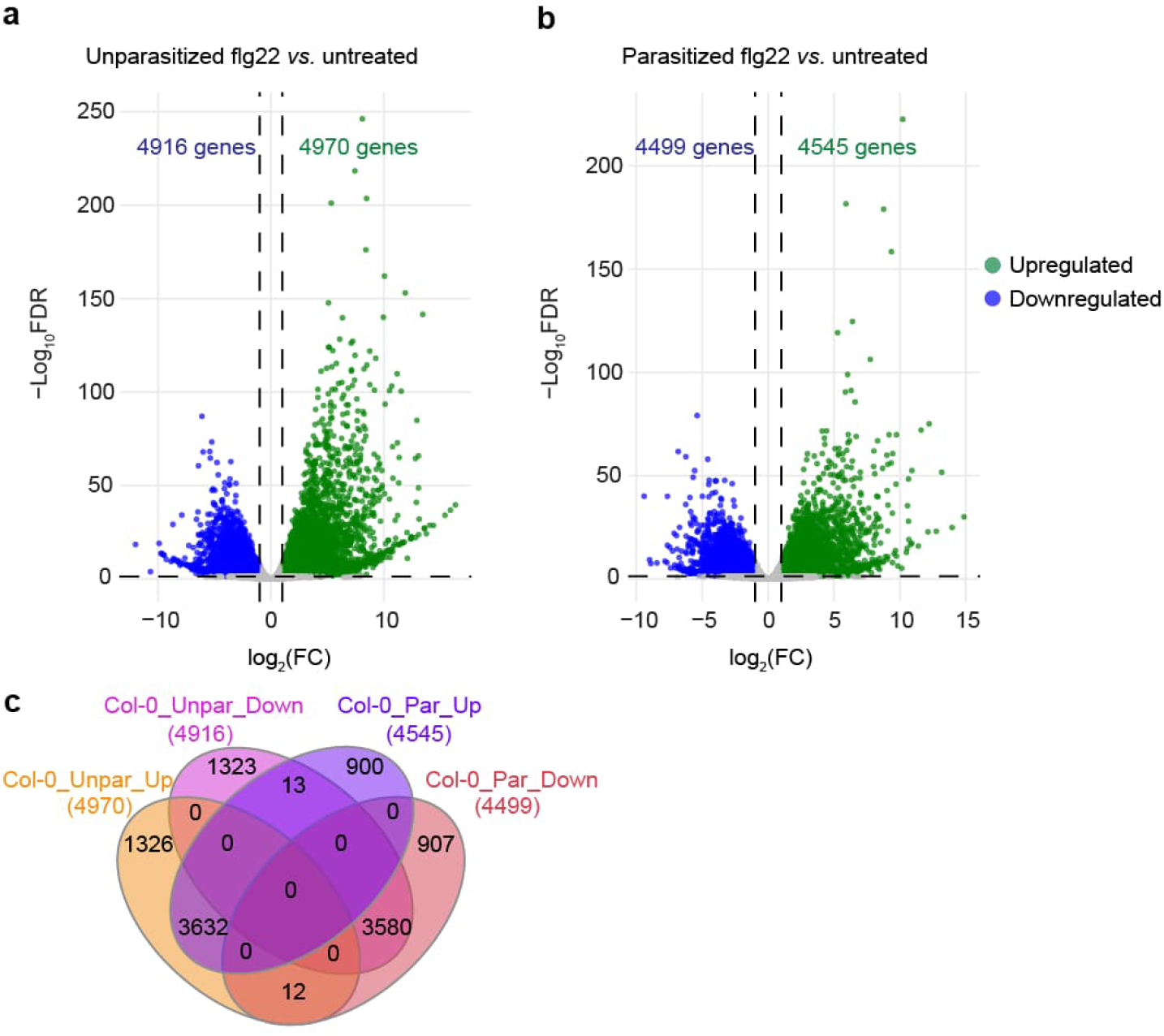
RNA-sequencing of unparasitized and parasitized *Arabidopsis* treated with flg22. **a, b**, Volcano plots showing the differentially expressed genes (DEGs) at 60 min after treatment with 100 nM flg22 in unparasitized *Arabidopsis* WT Col-0 plants relative to mock-treated controls **(a)** and in parasitized *Arabidopsis* WT Col-0 plants relative to mock-treated controls **(b)**. Green and blue points indicate significantly upregulated and downregulated genes, respectively (|log_2_FC| ≥ 1, FDR < 0.05). **c**, Venn diagram showing the unique and overlapping of upregulated and downregulated genes in unparasitized and parasitized *Arabidopsis* WT Col-0 plants after 24 hpi with *Pst* DC3000. Unpar, unparasitized; Par, parasitized.

**Extended Data Fig. 5.**
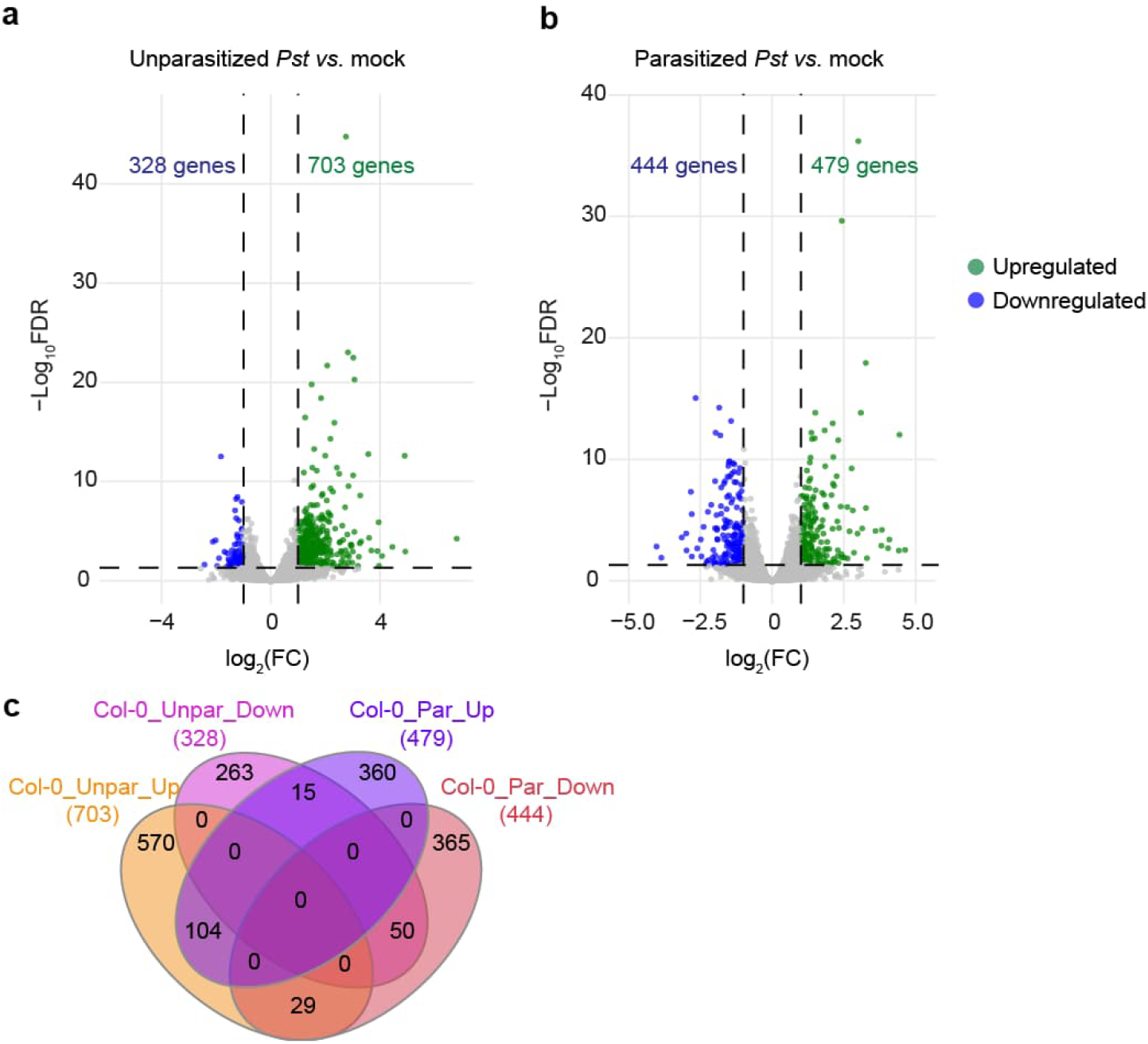
RNA-sequencing of unparasitized and parasitized *Arabidopsis* infected with *Pst* DC3000. **a, b**, Volcano plots showing the DEGs at 24 hpi with *Pst* DC3000 in unparasitized *Arabidopsis* WT Col-0 plants relative to untreated controls **(a)** and in parasitized *Arabidopsis* WT Col-0 plants relative to untreated controls **(b)**. Green and blue points indicate significantly upregulated and downregulated genes, respectively (|log_2_FC| ≥ 1, FDR < 0.05). **c**, Venn diagram showing the unique and overlapping of upregulated and downregulated genes in unparasitized and parasitized *Arabidopsis* WT Col-0 plants after treatment with 100 nM flg22. Unpar, unparasitized; Par, parasitized.

